## extended data for "Small-molecule P2RX7 activator sensitizes tumor to immunotherapy and vaccinates mouse against tumor re-challenge"

##### Generation of *p2rx7*$^{fl/fl}$ mice

P2rx7-flox mice were engineered as follow: ES clones (C57/BL/6) containing a construct for the conditional elimination or re-expression of P2RX7 (purchased from The European Conditional Mouse Mutagenesis Program, EUCOMM) were injected into blastocytes of C57Bl6/N, chimeric mice were selected and crossed with deleter mice that are transgenic for the Flip-recombinase under the control of the ubiquitous actin promoter to produce *p2rx7* $^{loxP/loxP}$ mice.

Our *p2rx7* $^{loxP/loxP}$ mice, with loxP sequence floxing the second exon of *p2rx7*, were crossed with (C57BL/6NTacGt(ROSA)26Sor<tm1(ACTB-Cre,-EGFP) transgenic mice which express the Cre-recombinase under the control of the b-actin promoter to produce p2rx7$^{exon2-/-}$ mice or with LysM-Cre mice (B6.129P2-Lys2tm1(cre)lfo from the Jackson Laboratories (obtained from Dr B. Chazaud, France) to generate myeloid cell conditional p2rx7 knockout and WT control (*p2rx7*$^{loxp/loxp}$) mice.

##### HEI3090 enhances the efficacy of anti-PD-1 antibody to inhibit melanoma growth in immunocompetent mice

To go further, we tested the efficacy of HEI3090 to inhibit the low immunogenic B16-F10 melanoma tumor growth. In the first setting of experiments, HEI3090 (1.5 mg/kg) or vehicle were administered daily for 11 days starting the day of tumor cell inoculations (Supplementary Fig. 3A). Mice treated with HEI3090 displayed significantly reduced tumor growth (left panel) and tumor weight (right panel). Flow cytometry analyses indicated that, as with LLC tumor cells, the success of HEI3090 treatment was associated with a better infiltration of immune cells within the tumor microenvironment (Supplementary Fig. 5A to D). The importance of mouse host cells to mediate HEI3090's antitumor effect was confirmed in experiment where *p2rx7*$^{-/-}$ mice were inoculated with B16-F10 cells. As described in the lung tumor model, HEI3090 was unable to inhibit melanoma growth when host cells were devoid of P2RX7 expression (Supplementary Fig. 4A). In the second setting of experiments, treatment started 3 days after tumor cell inoculation (at 2-8 mm$^2$ of tumor size). HEI3090 (3 mg/kg) was also able to inhibit tumor growth (Supplementary Fig. 3B), despite a lower efficacy than the one observed with LLC tumor cells (Fig. 2B). Of interest, HEI3090 treatment increases the median survival of 3 days (19 d. with HEI3090 vs 16 d. with vehicle). We also combined HEI3090 treatment with the anti-PD-1 immune checkpoint inhibitor and observed that HEI3090 enhanced the effect of the anti-PD-1 treatment. In particular, 2 out of 8 mice that were treated with HEI3090 combined to anti-PD-1 doubled their lifetime (Supplementary Fig. 3C).

**HEI3090 targets phagocytic cells, mainly dendritic cells, for its anti-tumor activity**

We have shown that HEI3090 requires immune cells to inhibit tumor growth (Fig 3A). It was of interest to see if HEI3090 targeted phagocytic cells to inhibit tumor growth, since they express high levels of P2RX7 and are able to kill tumor cells either directly or by inducing an immune response. To do so, we used several approaches. First, we depleted phagocytic cells by several administrations of clodronate liposome that has been described to deplete mainly macrophages and DCs. HEI3090's activity was lost in clodronate-liposome treated mice (Supplementary Fig4B) meaning that phagocytic cells are targeted by HEI3090. In order to specifically identify HEI3090-targeted cell, we conducted two complementary experiments using myeloid-specific P2RX7 deficiency (*p2rx7*$^{fl/fl}$-*LysM*) mice (Supplementary Fig4C) where P2RX7 was essentially lost in macrophages, and adoptive transfer of WT DCs into *p2rx7*$^{-/-}$ mice (Fig. 3B). HEI3090's anti-tumor activity was restored after adoptive transfer of WT DCs and was still efficient in *p2rx7*$^{fl/fl}$-*LysM* mice. The three experiments combined show that DCs are the major cell type targeted by HEI3090 to inhibit tumor growth. However, macrophages could still be involved in HEI3090-antitumor effect as a minor or secondary population but are not absolutely required like DCs.

We have also shown that IL-18 was necessary for HEI3090 anti-tumor effect (Fig 4). In order to see whether DCs produced IL-18, we quantified IL-18 in mice sera at the end of the WT DCs adoptive transfer experiment (D12). We can indeed see a slight increase in IL-18 release in sera of HEI3090-treated mice (Supplementary Fig9B). It is not that surprising not to see a major difference since mice received once WT DCs at the beginning of the experiment (D-1) and the IL-18 produced will be degraded since then. However, we can see that CD45$^{+}$ cells, CD8 and NK cells in the TME show an increase in the IFN-$\gamma$/IL-10 ratio in the HEI3090-treated mice (Supplementary Fig. 9B), showing that immune cells are differentiated into anti-tumor effector cells and suggesting that IL-18 produced by WT DCs was efficient during the experiment.

Collectively, these results suggest that the copresence of DCs, live tumor cells, ATP and HEI3090 represent the perfect microenvironment where DCs are primed by tumor antigens and activated by the ATP/P2RX7 axis to release IL-18 and inhibit tumor growth.

**HEI3090 does not favor immunogenic cell death**

P2RX7 expressed by DC has been shown to link innate and adaptive immune responses against dying tumor cells upon chemotherapy–induced ICD and facilitate tumor antigens presentation to T cells by the DC (6). In this study we choose two transplantable tumor models with different levels of P2RX7 expression. B16-F10 cells expressed higher level of P2RX7 than LLC cells (Supplementary Fig. 1B). To evaluate the capacity of HEI3090 treatment to induce ICD of tumor cells and concomitant stimulation of DC maturation, we first characterized whether HEI3090 induced tumor cell death. We observed that *in vitro* treatment of tumor cell lines (LLC and B16-F10) with BzATP (a stable agonist of P2RX7) and HEI3090 induced cell death *(*Supplementary Fig. 6A and B). We also noticed that HEI3090 treatment slightly increased P2RX7 expression in DC $CD4^{+}$ cells (Supplementary Fig. 4A). Second, we tested the HEI3090-induced ICD and DC stimulation hypothesis by inoculating WT mice with dying tumor cells and challenged them with live tumor cells, 7 days later. We used B16-F10 cells since they were more efficiently killed *in vitro* when stimulated by ATP and HEI3090 than LLC cells. Tumor growth was similar in mice vaccinated with *in vitro* treated tumors cells and in control mice (Supplementary Fig. 6C), indicating that the main antitumor effect of HEI3090 is not solely dependent on direct tumor ICD and the concomitant stimulation of DC maturation.

#### Supplementary Figure legends

##### Figure 1: Active P2RX7 is expressed on LLC and B16-F10 tumor cells

**A**. P2RX7 expression and activity in Lewis Lung carcinoma (LLC) cell line. P2RX7 expression was assayed by flow cytometry using the anti P2RX7 (clone 1F11, left panel) or by western blotting using an anti-extracellular loop of P2RX7 antibody (APR-008, Alomone Labs) that recognizes a band of 70kDa, the expected size for P2RX7 (middle panel). To assay P2RX7's activity we measured the ATP-induced increase of intracellular $Ca^{2+}$ concentration (Fluo-4-AM uptake) in the presence of the P2RX7's inhibitor (GSK1370319A), using a plate reader (right panel). **B**. P2RX7 expression and activity in melanoma (B16-F10) cell line. These results demonstrated that both cell lines expressed an active P2RX7, with B16-F10 cells expressing higher P2RX7 levels than LLC cells. Bars are mean +/- SEM; *<0.05; **<0.01; ***<0.001 (t-test).

##### Figure 2: Picture of HEI3090 effect on LLC tumor size

$5x10^5$ LLC were injected s.c. into the flank of C57BL6/J mice. Mice were injected i.p every day with vehicle or HEI3090 (1.5 mg/kg) from day 1 to day 12. Pictures showed tumor the day of the sacrifice. Bar = 10 mm

##### Figure 3: HEI3090 inhibits the growth of melanoma B16-F10 tumor cells and increased mice survival when combined with anti-PD-1 antibody

**A.** $5x10^5$ B16-F10 were injected s.c. into the flank of C57BL6/J mice. Mice were injected i.p every day with vehicle or HEI3090 (1.5 mg/kg) from day 1 to day 12. Curves showed mean tumor area in $mm^2$ (left panel) and graph showed mean tumor weight the day of sacrifice (right panel). **B.** Effect of HEI3090 in therapeutic administration. Mice were treated i.p. with vehicle or HEI3090 (3 mg/kg) from days 5 (when tumors reached a volume of approximatively 5-10 $mm^2$) to 15. Curves showed mean tumor area (left panel) and survival of B16-F10 tumor bearing mice (right panel). **C.** Effect of HEI3090 and $\alpha$PD-1 checkpoint inhibitor on survival of B16-F10 tumor bearing mice. $5x10^5$ B16-F10 were injected s.c. into the flank of WT mice. Mice were treated i.p. with vehicle or with HEI3090 (3 mg/kg) from days 1 to day 18. Each mouse received 200 µg of $\alpha$PD-1 i.p. at day 4, 7, 10, 12 and 16. Mice were sacrificed when tumor reached 100 $mm^2$. Bars are mean ± SEM. *p<0.05, **p<0.01 ***p<0.001, (Mann-Whitney or Mantel Cox (C) tests) Each point represents one mouse. **D**. Efficacy of the combo treatment in the *in situ* (LSL $KRas^{G12D}$) genetic lung tumor mouse model. Ten lung lesions per mouse (5 large, 5 small) were selected. Average cell number/$mm^2$ and $Ki67^+$ cells are shown. Bars are mean ± SEM. *p<0.05,

**$p<0.01$ ***$p<0.001$, ****$p<0.0001$ (Mann-Whitney or Mantel Cox (C and E) tests). Each point represents one individual mouse.

**Figure 4: HEI3090 targets P2RX7-expressing dendritic cells.**

**A.** P2RX7 expression by DC $CD4^+$ cells was increased in response to HEI3090. **B.** Three days prior to tumor cell inoculation, liposome clodronate (200 µl) was injected ip to mice. Then mice were injected every 3 days with liposome clodronate 1h before HEI3090 treatment. $5.10^5$ LLC cells were inoculated s.c. and mice were treated daily with 1.5 mg/kg of HEI3090 or vehicle. Tumor area was measured with a caliper. At the end of the experiment, tumors were weighted. **C.** Cells from the myeloid lineage did not support HEI3090-induced antitumor response. $5x10^5$ LLC cells were injected s.c. to *p2rx*$7^{fl/fl}$ -LysM mice and mice were daily treated with HEI3090 or vehicle, as detailed in the method section. Curves showed mean tumor area in $mm^2$ (left panel) and graph showed mean tumor weight the day of sacrifice (right panel). Each point represents one individual mouse ***$p<0.001$(Mann-Whitney test).

**Figure 5: Immune cells mediate the antitumor activity induced by HEI3090 in the B16-F10 tumor mouse model**

**A.** Effect of HEI3090 on B16-F10 tumor growth in *p2rx7*$^{-/-}$mice. $5x10^5$ cells were injected s.c. into the flank of *p2rx7*$^{-/-}$ mice. Mice were treated i.p. with vehicle or with HEI3090 as indicated in A. Curves represent daily follow up of tumor area (left panel) and graph corresponds to the weight of tumors the day of sacrifice (right panel). **B.** Effect of HEI3090 on the recruitment of immune cells within the TME. $5x10^5$ B16-F10 cells were injected s.c. into the flank of WT mice. Mice were injected i.p every day with vehicle or HEI3090 (1.5 mg/kg). At day 12, tumors were collected for flow cytometry analyses. **C.** Proportion of M-MDSC within the TME among $CD45^+$ within LLC tumors from mice treated with vehicle or HEI3090. In this tumor model PMN-MDSC were not affected by HEI3090 treatment (not shown). **D.** Ratio of NK, $CD4^+$ or $CD8^+$ T cells on M-MDSC within the TME. *$p<0.05$, **$p<0.01$ ***$p<0.001$, ****$p<0.0001$ (Mann-Whitney or two-way Anova (C) tests). Each point represents one individual mouse.

**Figure 6: HEI3090 does not promote immunogenic cell death**

HEI3090-induced LLC (**A**) and B16-F10 (**B**) tumor cell toxicity as described above. **C.** *In vivo* assay for the evaluation of HEI3090 as an inducer of immunogenic cell death. B16-F10 were exposed to 3mM ATP and 50µM HEI3090. WT mice were inoculated s.c. with $1.10^5$ dying B16-F10 cells in the right flank (n=6) or PBS as control (n=4). 7 days later, $5.10^5$ live B16-F10 cells were injected s.c. in the contralateral flank. The details of the ICD experiment are provided in the methods section. Curves shown mean of tumor growth in the contralateral site. Bars are mean ± SEM. *p<0.05, **p<0.01 ***p<0.001, (Mann-Whitney test).

**Figure 7: HEI3090 increased eATP-induced caspase-1 cleavage in macrophages**

$4.10^5$ peritoneal macrophages from WT mice were primed for 4h at 37°C with 100 ng/ml LPS and then stimulated for 30 minutes with 10 µM nigericin or 3 mM ATP with 50 µM HEI3090 or DMSO. Whole cell protein extracts were analyzed by western blotting by using antibodies recognizing NLRP3, ASC, pro-caspase-1 and active form caspase-1, and β-actin as a loading control. Relative band intensities of each protein were assessed to that of β-actin. Bar are mean +/- SEM. Each point represents one individual experiment. *<0.05 (t-test).

**Figure 8: HEI3090-induced IL18 production required P2RX7 expression**

**A.** IL-18 staining of LLC tumors collected from WT, *p2rx7*$^{-/-}$ or *il18*$^{-/-}$ mice, demonstrated the specificity of IL-18 antibody **B.** Production of IL-18 (ELISA) by splenocytes isolated from *p2rx7*$^{-/-}$ mice, demonstrated that without P2RX7 expression, HEI3090 was unable to increase IL-18 levels. **C.** IL-18 staining from lung of LSL *KRas*$^{G12D}$, illustrating that IL-18 expression was increased in lung macrophages of HEI3090 + αPD-1 treated mice. **D.** Production of IL-1β (ELISA) by peritoneal macrophages isolated from WT mice, demonstrated that HEI3090 was unable to increase IL-1β levels. The specific inhibitor of NLRP3 (MCC950 compound) efficiently inhibits IL-1β production. **E.** Quantification of IL-1β levels from serum of indicated mice confirming that HEI3090 did not modulate the production of IL-1β in *p2rx7*$^{-/-}$ complemented with WT DC. **F**. Quantification of IL-1β levels from serum of LSL *KRas*$^{G12D}$ mice treated with HEI3090 and anti PD-1 antibody. Each point represents one mouse. Bars are mean ± SEM. *p<0.05, **p<0.01 (Mann-Whitney or two-way anova (B, C and D) tests.

**Figure 9: Indirect effect of HEI3090 on IFN-γ production by antitumor immune cells**

**A.** *Ex vivo* IFN-γ production. Splenocytes from WT mice were treated as indicated and IFN-γ production was assayed by flow cytometry in NK, $CD4^+$ and $CD8^+$ T cells. **B.** Purified WT DC were inoculated to *p2rx7*$^{-/-}$ mice i.v. at D-1. At D0, $5x10^5$ LLC cells were injected s.c.. Mice were treated i.p. with vehicle or with HEI3090 daily for 11 days. At D12, mice were sacrificed, sera and tumors were collected for ELISA and flow cytometry analyses respectively. The ratio of IFN-γ on IL-10 in indicated cells (left panel) and the concentration of IL-18 in the sera (right panel) are shown. **C.** MHC-I and PD-L1 expression induced by HEI3090 on LLC *in vitro*. LLC were stimulated as indicated for 24h and expression of MHC-I ($H2K^d/D^b$)

and PD-L1 were assayed by flow cytometry analyses. This data revealed that P2RX7 activation indirectly led to increased tumor immunogenicity.

**Supplementary Figure 1**

Lung tumor cells (LLC)

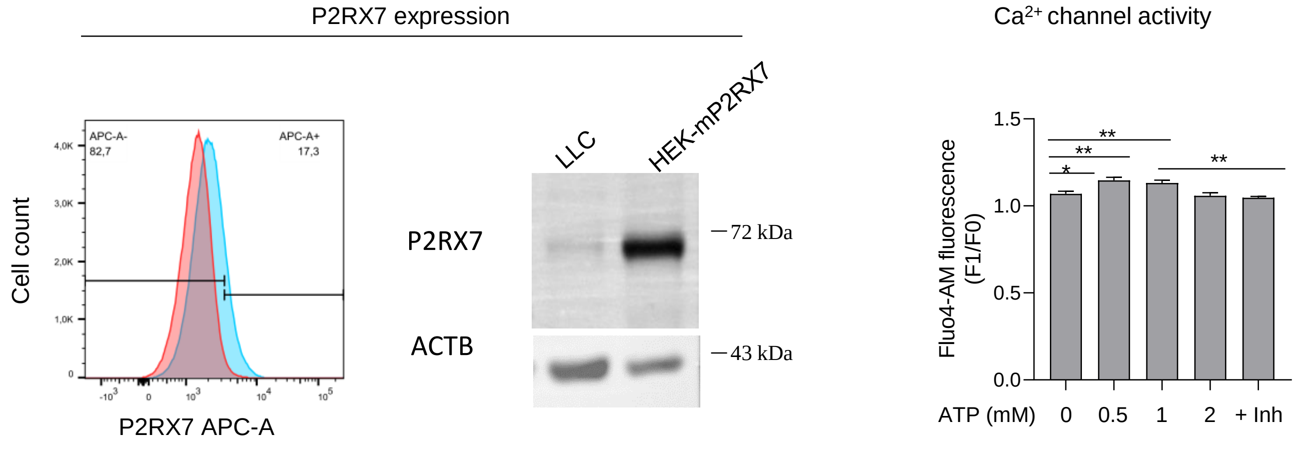


Melanoma tumor cells (B16-F10)

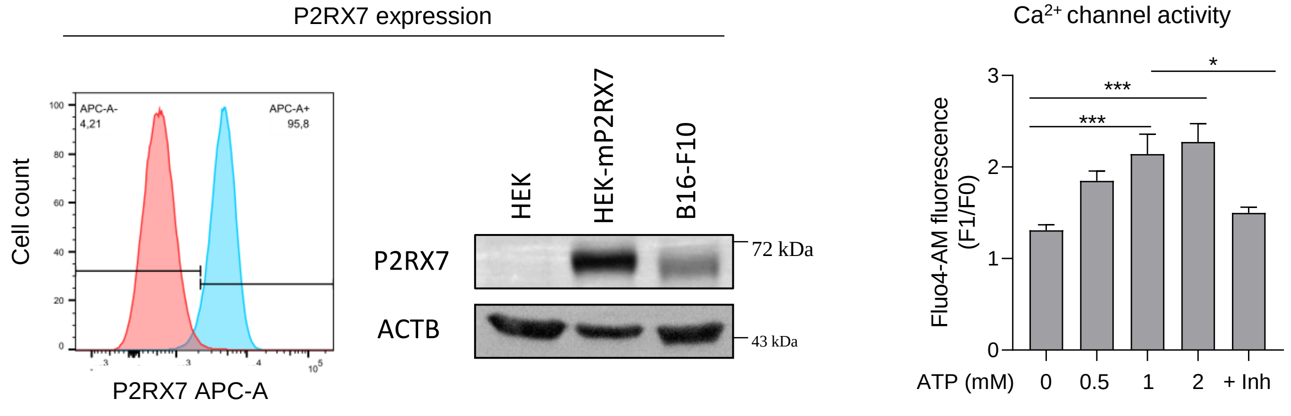

#### Supplementary Figure 2

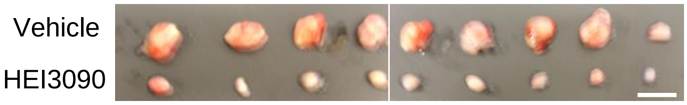


#### Supplementary Figure 3

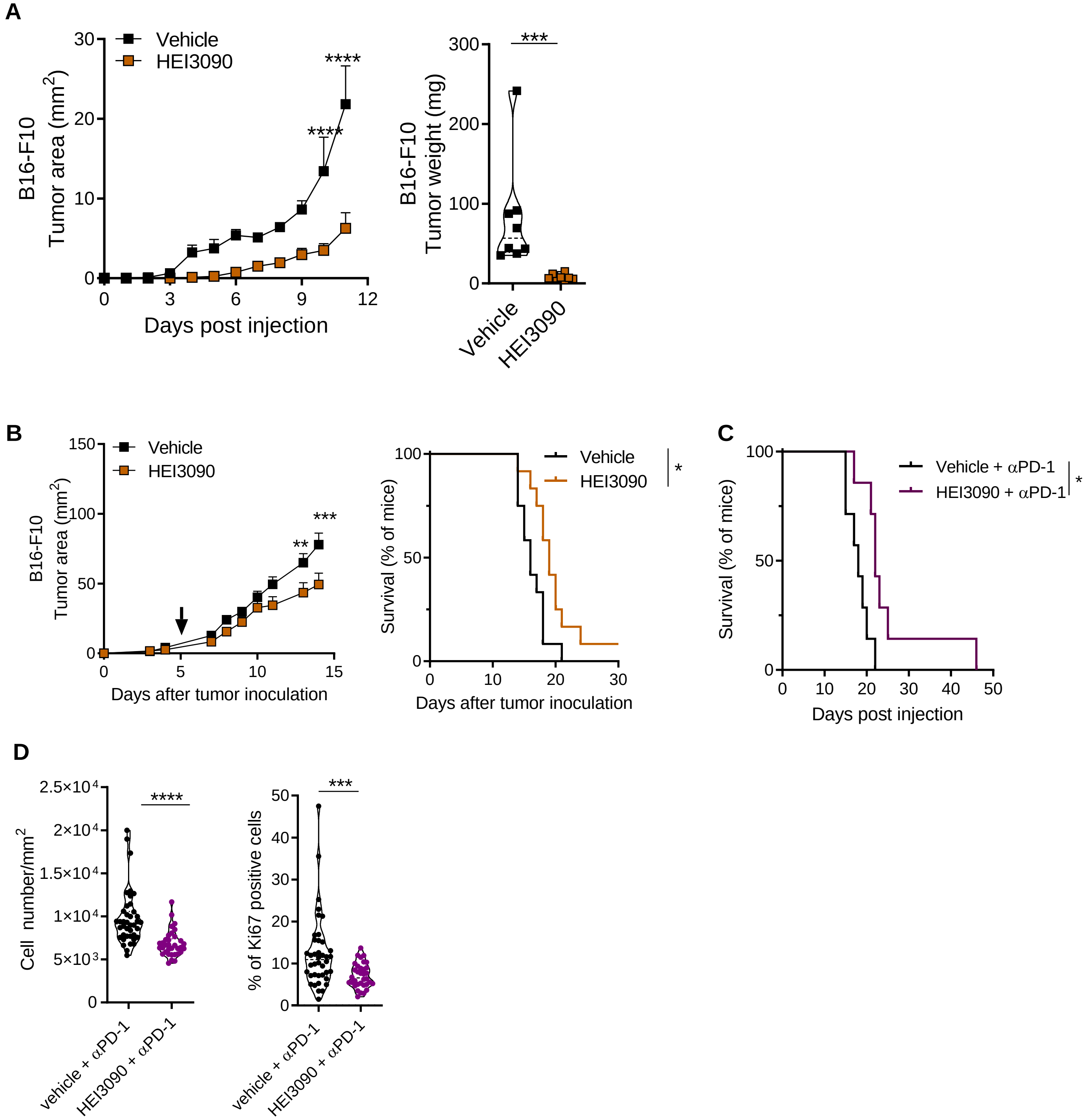

**Supplementary Figure 4**

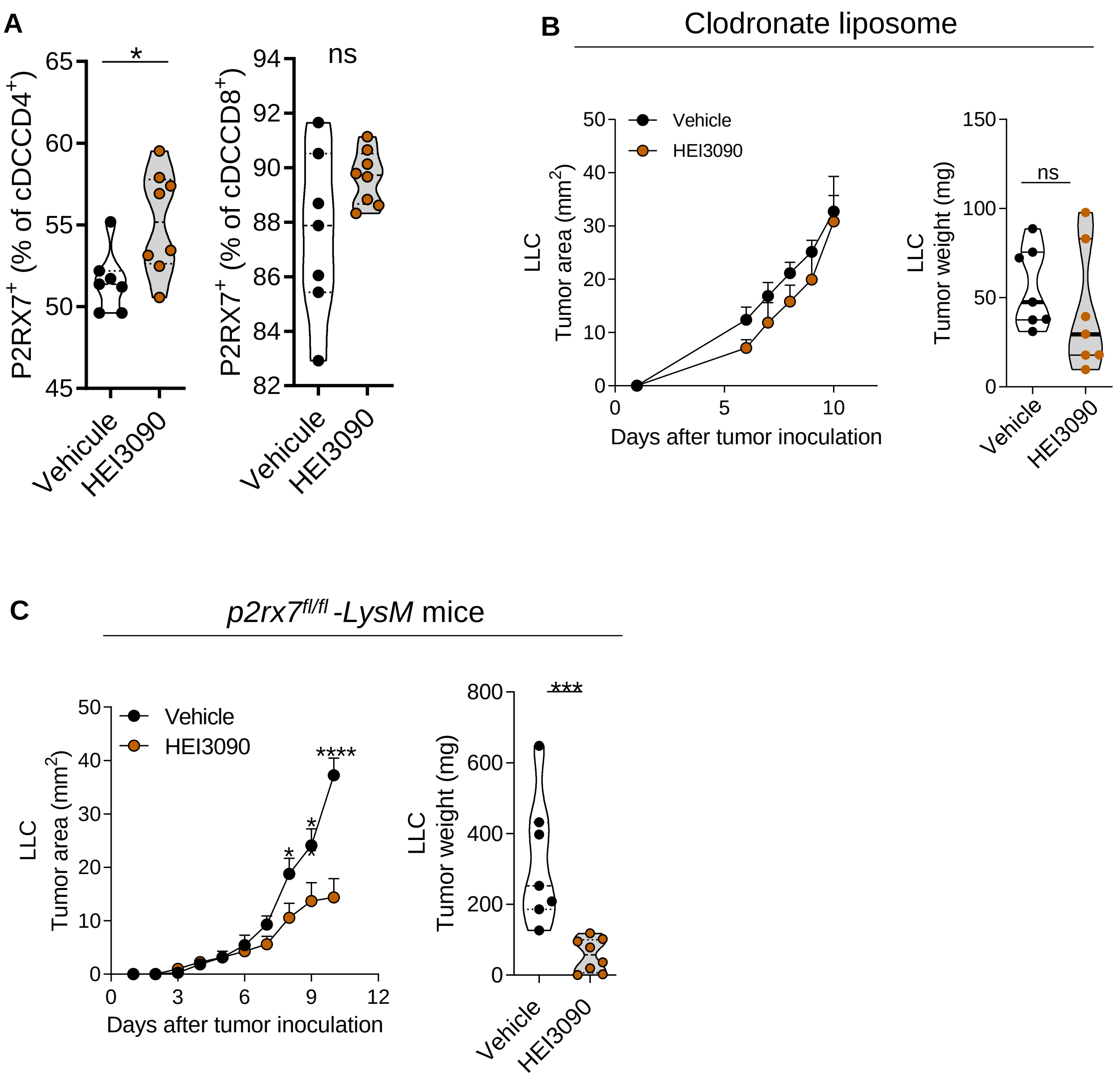

#### Supplementary Figure 5

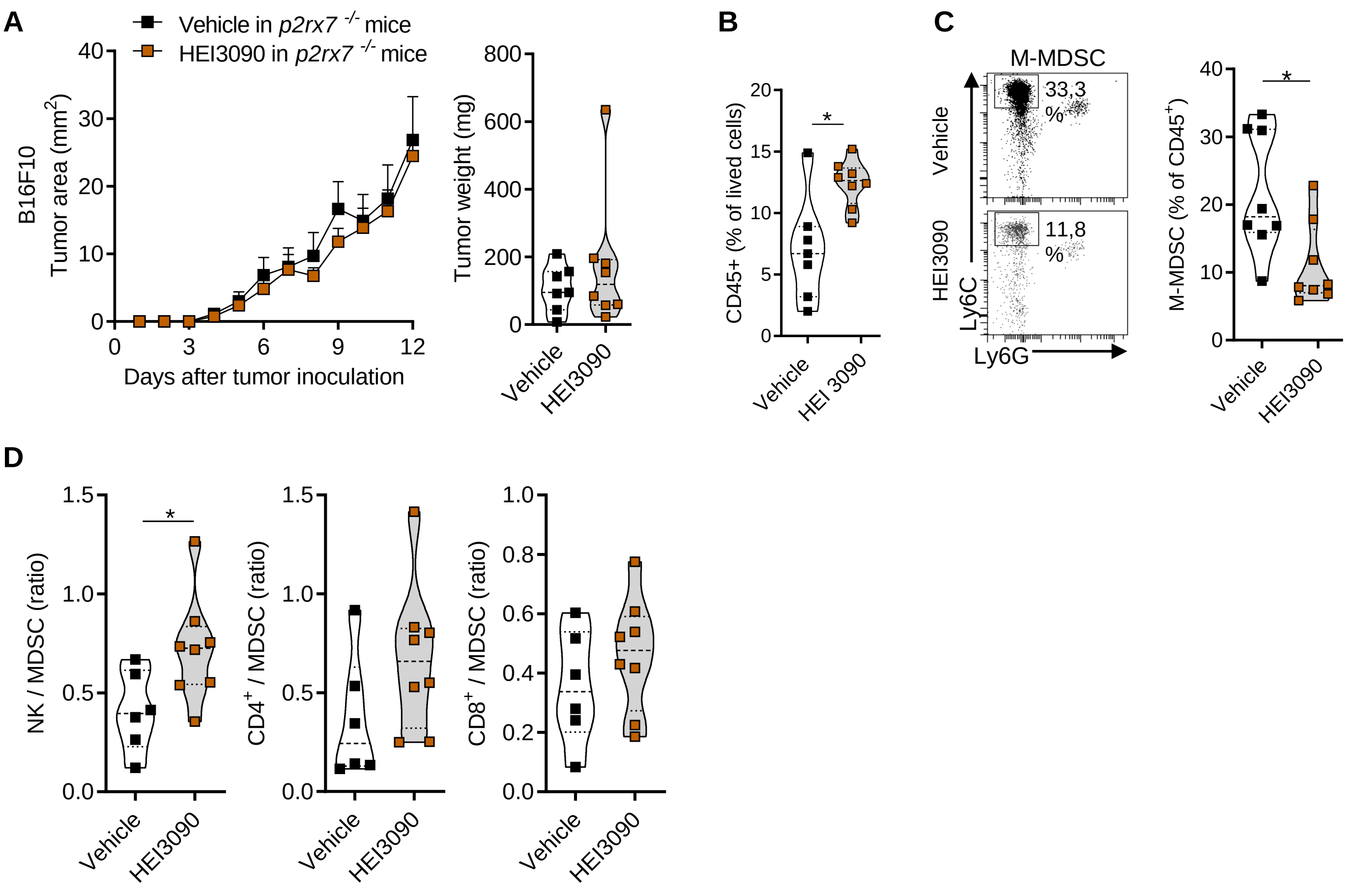

#### Supplementary Figure 6

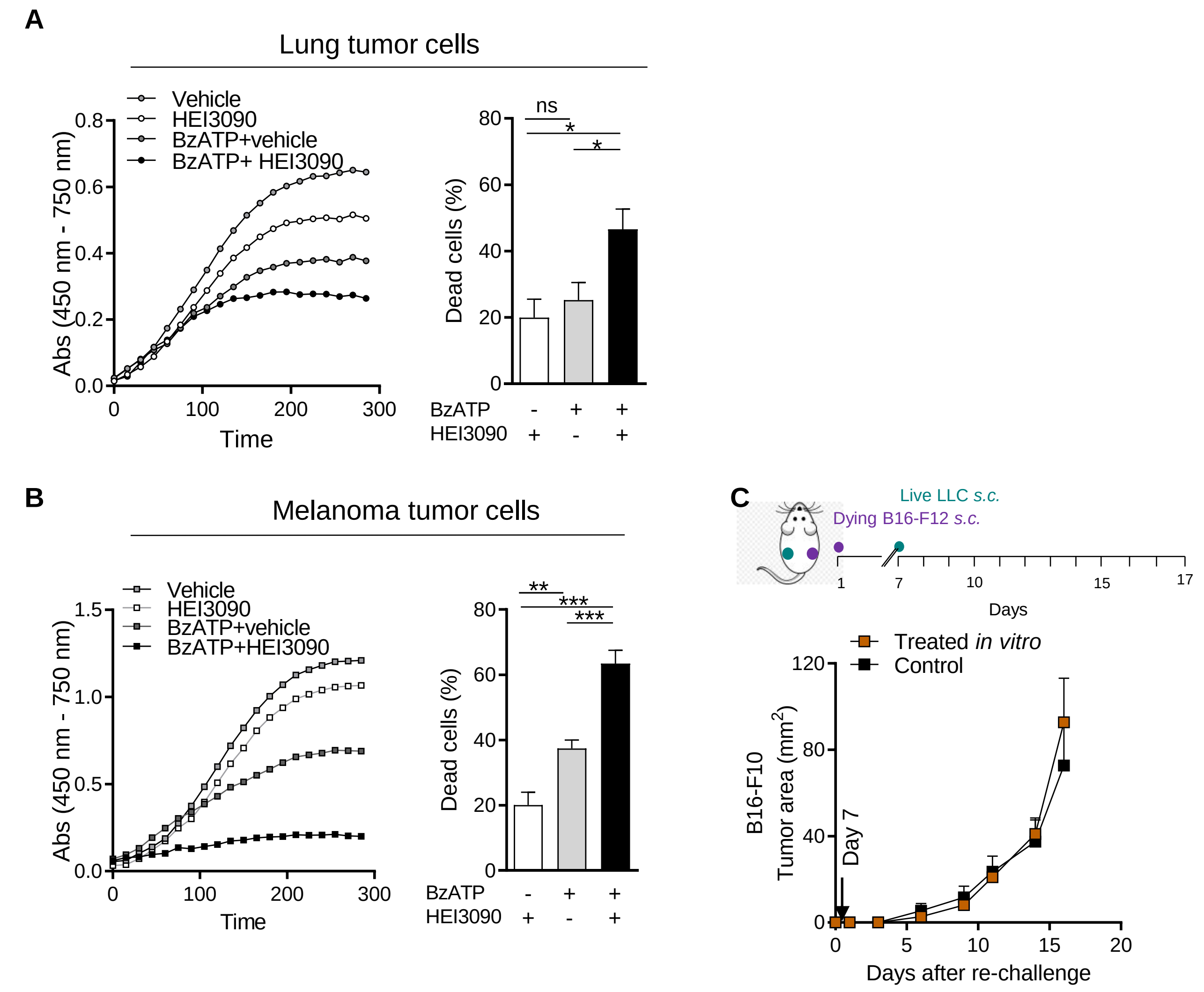


#### Supplementary Figure 7

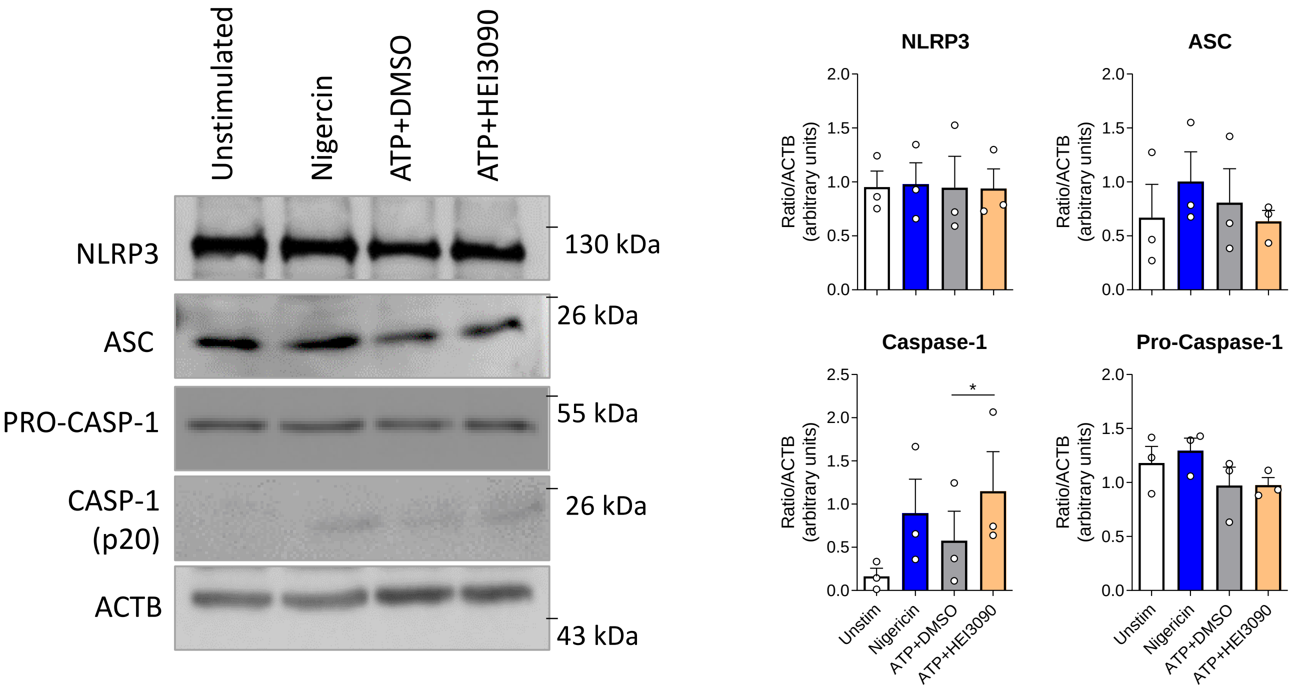

### Supplementary Figure 8

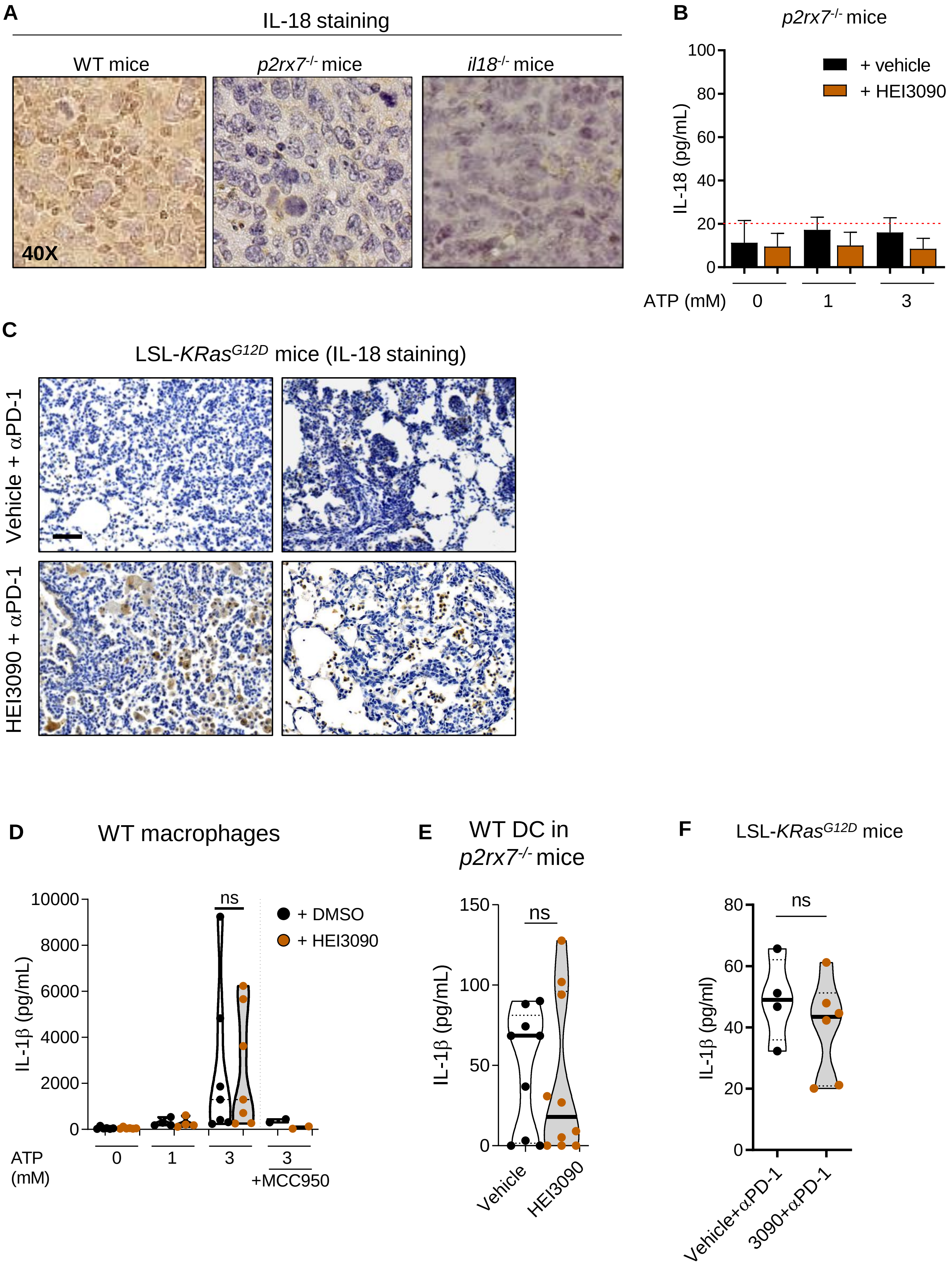

#### Supplementary Figure 9

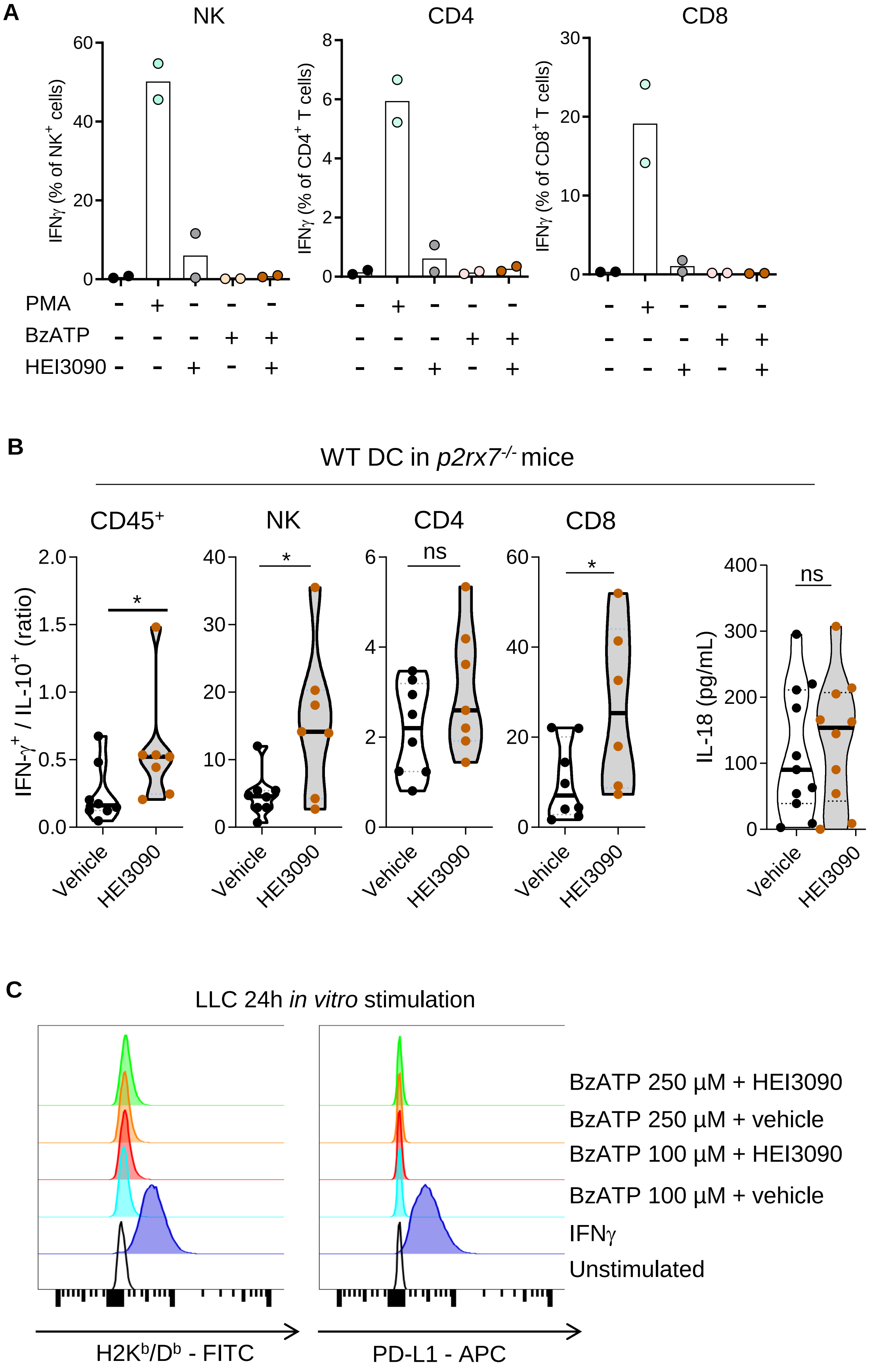
